# Information Enhanced Model Selection for Gaussian Graphical Model with Application to Metabolomic Data

**DOI:** 10.1101/815423

**Authors:** Jie Zhou, Anne G. Hoen, Susan McRitchie, Wimal Pathmasiri, Weston D. Viles, Quang P. Nguyen, Juliette C. Madan, Erika Dade, Margaret R. Karagas, Jiang Gui

**Author notes:** Corresponding author, *Email address:* (Jiang Gui).

## Abstract

In light of the low signal-to-noise nature of many large biological data sets, we propose a novel method to learn the structure of association networks using Gaussian graphical models combined with prior knowledge. Our strategy includes two parts. In the first part, we propose a model selection criterion called structural Bayesian information criterion (SBIC), in which the prior structure is modeled and incorporated into Bayesian information criterion (BIC). It is shown that the popular extended BIC (EBIC) is a special case of SBIC. In the second part, we propose a two-step algorithm to construct the candidate model pool. The algorithm is data-driven and the prior structure is embedded into the candidate model automatically. Theoretical investigation shows that under some mild conditions SBIC is a consistent model selection criterion for high-dimensional Gaussian graphical model. Simulation studies validate the superiority of the proposed algorithm over the existing ones and show the robustness to the model misspecification. Application to relative concentration data from infant feces collected from subjects enrolled in a large molecular epidemiological cohort study validates that metabolic pathway involvement is a statistically significant factor for the conditional dependence between metabolites. Furthermore, new relationships among metabolites are discovered which can not be identified by the conventional methods of pathway analysis. Some of them have been widely recognized in biological literature.

## 1. Introduction

Modern ‘omics technology can easily generate thousands of measurements in a single run which provides an opportunity for researchers to explore complex relationships in biology. However, it has been widely recognized that biological measurements are usually accompanied by a low signal-to-noise ratio making detection of effect challenging and final conclusions unreliable. As reported in Ideker et al [31], prior knowledge can play a pivotal role in deciphering this kind of complexity. For example, Segre et al [67] drew on the prior knowledge on mitochondrial genes sets to investigate whether mitochondrial dysfunction is a cause of the common form of diabetes. Roach et al [64] identified the gene that causes Miller syndrome based on the human genome reference map. For more work on the application of prior biological knowledge, see Boluki et al [7], Imoto et al [32], Ma et al [48]. The studies in this paper are motivated by the metabolite pathway information that are available in many public biological databases such as Kyoto Encyclopedia of Genes and Genomes (KEGG). As far as we know, such information have not been well utilized in literature to improve the statistical analysis of metabolites.

Biological network, such as microbe-microbe interaction networks, metabolite association networks and gene regulation networks, have received much attention in recent years. Probabilistic graphical models are typically employed in literature to study such biological networks (Lauritzen and Sheehan [42], Zhang et al [86], Stingo et al [72], Dobra et al [19]). The edges in graphical model represent the conditional dependence among the vertices, which stand for the objects of interest such as microbes, metabolites or genes. In some cases, i.e., causal inference, the directionality of edge, which is indispensable for the computation of casual effects, also has to be considered. Nevertheless, in this paper we only consider association network which can be characterized by undirected graphical model. The identification of the structure of association network is our primary interest. Many algorithms have been proposed in literature in this respect. In particular, for tree and forest, Chow and Liu [17], Edwards et al [21] and Kirshner et al [38] proposed and studied the classic Chow-Liu algorithm and its extensions. Heuristic algorithms such as the hill-climbing algorithm are studied in Hojsgaard et al [30], Jalali et al [36], Lauritzen [41] and Ray et al [62]. Cheng et al [15], Friedman et al [22], Meinshansen and Buhlmann [49], Ravikumar et al [61] and Wainwright and Jordan [79] investigated the *L*_1_-penalized likelihood method for the identification of Gaussian and Ising graphical models.

In order to deal with the prior structure of network, Bayesian methods is the typical choice in literature (Dobra et al [19], Jones et al [35], Scott and Berger [66]). As for Gaussian graphical model (GGM), the most popular Bayesian method is based on the direct modeling of prior distribution of precision matrix, e.g, conjugate G-wishart distribution (Mohammadi and Wit [52]), or spikeslab distribution (Mohammadi [53]) et al. In these situations, though there exist MCMC sampling algorithms for the decomposable graph, the computation becomes challenging for the general non-decomposable graph (Carvalho et al [11], Dobra and Lenkoski [20], Mitsakakis et al [51], Roverato [65], Wang [80], Wang and Carvalho [81]). Recently, an efficient sampling-free Bayesian method is proposed in Leday and Richardson [43] for high-dimensional GGM, which employs hypothesis testing to determine the existence of each individual edge based on Bayes factor. There are also several algorithms to deal with the prior information within frequentist framework. For example, the Chow-Liu algorithm can learn the structure when the graph is a tree (Chow and Liu [17]); Chow-Liu algorithm is extended in Edwards et al [21] to deal with the forest; conditional Chow-Liu algorithm is proposed in Kirshner et al [38] to include a given set of edges in the graph. In extended Bayesian information criterion (EBIC) for high-dimensional Gaussian graphical model (Foygel and Drton [26]), as will be shown in Section 3, empty graph is adopted as the prior graph. If a sufficient large weight is assigned to the prior structure, the algorithm will end up with an empty graph. Ma et al [48] focus on the deterministic prior structure; when the random structure is involved in prior graph, they propose an algorithm to construct the model pool.

In this paper we propose a novel method to learn the structure of Gaussian graphical models based on a given prior structure. We first propose a model selection criterion called structural Bayesian information criterion (SBIC) to incorporates the prior structure into BIC. Numerous criteria have been proposed in literature for model selection, e.g., Akaike information criterion (AIC, Akaike [1]), Bayesian information criterion (BIC, Schwarz [69]), extended BIC (EBIC, Bogdan et al [4, 5, 6], Chen and Chen [13, 14]), cross-validation method (CV, Stone [75]), generalized CV method (GCV, Geisser [28], Burman [10], Shao [70], Zhang [83]), risk inflation criterion (RIC, Foster and George [25], Zhang and Shen [85]) among many others. In particular, generalized information criterion (GIC) proposed in Shao [71], Kim et al [37] provided a unified framework for many of these criteria in the context of linear regression model, such as AIC, BIC, EBIC and RIC et al. As for the network selection, Foygel and Drton [26] studied the consistency of EBIC for GGM selection. The criterion SBIC proposed in this paper can be regarded as a generalization of the EBIC of Foygel and Drton [26]. Compared with the EBIC, SBIC provides a more flexible framework; in fact, SBIC just reduces to EBIC when the prior graph is an empty graph. As a theoretical basis, for high-dimensional sparse Gaussian graphical models, it is shown that SBIC is a consistent criterion for model selection under mild conditions. Based on the prior structure, we then propose a data-driven two-step algorithm to build the model pool, in which the graph is enriched in the first step and pruned in the second step. Such two-step algorithm can be readily implemented based on the R packages such as *glmnet* (Friedman et al [22]) or *glasso* (Friedman et al [24]). Recall that the well-known greedy equivalence search (GES) algorithm for structure learning of directed acyclic graph also consists of similar edge addition and removal steps aiming to optimize the score function. For decomposable graph, GES algorithm converges to the global optimum in probability as *n* → ∞ (Chickering et al [16]).

Through simulation studies, it is shown that the combination of the proposed SBIC and two-step algorithm is a robust structure-learning strategy for highdimensional Gaussian graphical model and outperforms many existing popular structure learning algorithms under the given conditions. As an application, we studied ^1^H NMR-based metabolite data profiled in infant feces collected as part of the New Hampshire Birth Cohort Study (NHBCS), a large prospective cohort study of mothers and their children born in New Hampshire (Madan et al [46]). The prior structure for these metabolites is constructed based on the related pathway information from KEGG. Our results show that pathway is a statistically significant factor for the conditional dependence between metabolites, i.e., the strength of conditional dependence between two metabolites increases if the proportion of shared pathways increases. Furthermore, the identified network discovers new relationship between metabolites that can not be identified through the conventional methods of pathway analysis, many of which have been validated in biological literature.

The paper is organized as follows. Section 2 briefly reviews the undirected Gaussian graphical model and extended BIC. A new formulation of extended BIC is introduced. In Section 3, we present our main algorithm, in which Section 3.1 introduces structural BIC and its implications; Section 3.2 describes the two-step algorithm for building the candidate model pool. Theoretical investigation of SBIC is given in Section 4. In Section 5, the algorithm is evaluated through simulated data. In Section 6 we use the algorithm to investigate the pathway and metabolomic data from NHBCS. Section 7 concludes with a brief comment.

## 2. Gaussian Graphical Model and EBIC

### 2.1. A brief review of EBIC for Gaussian graphical model

For a givenp-dimensional multi-normal random vector, **X** = (*X*_1_, *X*_2_, ···, *X_p_*)^*T*^ ~ *N*(***μ***_*p*_, Σ_*p*×*p*_), an undirected graph *G* = (*V, E*) is used to represent the conditional dependence relationship between **X**, where the vertex set *V* indexes the variables and the edge set *E* encodes the conditional independence. The precision matrix is defined as Ω_*p*×*p*_ = (*ω_ij_*) = Σ^-1^. It turns out that for multi-normal distribution the precision matrix can completely specify the structure of *G*. Given *n* i.i.d. observations, 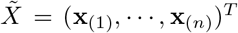, our aim is to learn the structure of *G*, i.e., to identify the nonzero components in 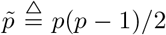 off-diagonal entries in Ω. In its general form, Bayesian information criterion (BIC) can be stated as follows. Let 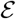 be the model space under consideration with *π*(*E*) the prior probability for 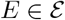. Let *θ* denote the unknown parameter in *E* with prior distribution *p*(*θ*). With *θ* in hand, let the conditional density function for 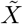 be 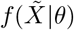, then the marginal density function for observations 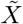 can be expressed as 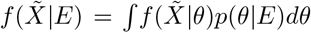. Therefore, the posterior distribution of model *E* can be expressed as

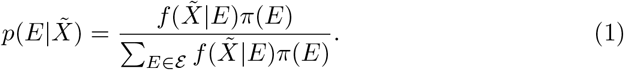

Through Laplace’s method of integration, the following approximation can be obtained for 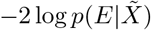,

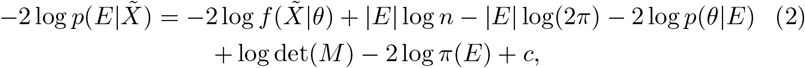

where *M* is the expected information matrix for single observation, |*E*| the degree of freedom of model *E* and *c* a constant. By omitting the last five terms which do not involve the sample size *n*, we get the standard Bayesian information criterion, BIC(*E*) = – 2*l_n_*(*E*) + |*E*| logn with 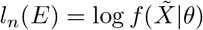. For the highdimensional regression model, the extended BIC (EBIC) is proposed in Bogdan et al [4], Bogdan et al [5], Bogdan et al [6], Chen and Chen [13], Chen and Chen [14], which puts more weight on sparse model than standard BIC. EBIC is further generalized to the Gaussian graphical model in Foygel and Drton [26], which has the following form,

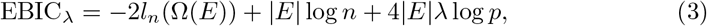

where Ω(*E*) is the precision matrix associated with model *E*. Tuning parameter 0 ≤ λ ≤ 1 controls the model complexity. When λ = 0, EBIC reduces to the standard BIC. As λ increases, (3) will put more weight on the sparse model. The log-likelihood function *l_n_*(Ω(*E*)) in (3) for the Gaussian graphical model has the following form,

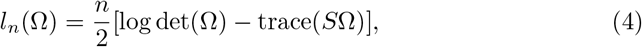

where *S* is the empirical covariance matrix. It is proved in Foygel and Drton [26] that EBIC_λ_ (3) is a consistent model selection criterion for high-dimensional GGM under some mild conditions.

Although EBIC has been widely used in literature for high-dimensional model selection, several limitations have not been addressed adequately. In particular, EBIC does not take prior information into consideration. Typically, prior information is integrated with data through Bayesian method; however, as we have mentioned in previous section, the complicated forms of posterior distribution for non-decomposable graphs have undermined the popularity of Bayesian method in practice. In this context, incorporating prior information into BIC is a natural choice. On the other hand, given such model selection criterion, an appropriate candidate model pool is indispensable since the exhaustive search within the whole model space is impossible for high-dimensional graphical model. With these motivations in mind, in the following section we propose a new strategy for the selection of Gaussian graphical model when the prior structure is available. Though we focus on Gaussian graphical model in this paper, the algorithm can be easily adapted to accommodate more general undirected graphical model, e.g., Ising model.

### 2.2. A new interpretation of EBIC

In this section, we introduce a different way to interpret EBIC which will facilitate the introduction of prior structure in Section 3. For any given pair of vertices, (*X_i_, X_j_*), define the edge variable *Z_ij_* equal to one if there exists an edge between *X_i_* and *X_j_* and zero otherwise, i.e., *Z_ij_* is the indicator variable for the existence of the edge between nodes *X_i_* and *X_j_*. Due to the symmetry of undirected graph, we have *Z_ij_* = *Z_ji_* (1 ≤ *i* < *j* ≤ *p*). Then we define a 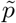-dimensional random vector 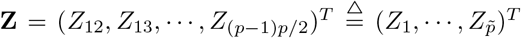. The prior information about the structure of *E* can be completely specified by the probability distribution of **Z**. The following Boltzmann distribution is employed to model the distribution of **Z**,

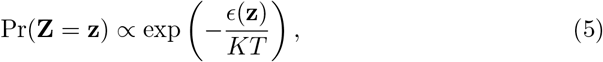

where *ϵ*(·) ≥ 0 is called energy function, *T* the temperature parameter, and *K* the Boltzmann constant. Note that there is a one-to-one correspondence between **Z** = **z** and model *E* so that (5) amounts to defining a prior distribution *π*(*E*) for *E*. Without loss of generality, *K* = 2 is always assumed in the following discussion. Substitution of (5) into (2) yields the following approximation to (2) for Gaussian graphical model,

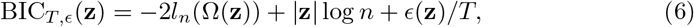

where |**z**| denotes the number of nonzero components in **z**. The third to fifth terms and the constant *c* in (2) have been omitted here since neither sample size *n* nor model dimensionality *p* is involved in these terms. In order to use (6) in practice, we consider the following simple yet flexible quadratic specification of *ϵ*(·),

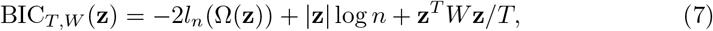

for some given positive semi-definite matrix W. In (7), the energy function can be regarded as the squared weighted Euclidean distance between the given state, **z**, and the origin state, **0**. It is obvious that both BIC and EBIC are the special cases of (7). In fact, if *W* = 0, (7) is the standard BIC; if *T* = 1/(4λ) and 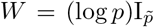 with 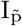 the 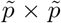 identity matrix, then (7) reduces to the EBIC (3)–(4). With such a specification of *W* in EBIC, it is straightforward to show that the components of **Z** are independent Bernoulli variables with nonzero probability 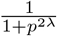. Such probabilistic interpretation can guide us to choose a proper tuning parameter λ involved in EBIC (3). For example, for λ = 0.5, or equivalently *T* = 0.5, which is often recommended in literature (Foygel and Drton [26]), it implies that the prior mean number of total edges is 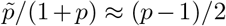. More generally, we have *P*(*Z_i_* = 1) < 0.5 whenever *T* > 0 and *P*(*Z_i_*) > 0.5 whenever *T* < 0. So for the sparse graph with 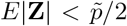, *T* > 0 is a plausible choice.

In some circumstances, prior information comprise both the mean and variance of the total edges, which can also be modeled through BIC_*T,W*_. Specifically, consider the following form of *W* for BIC_*T,W*_,

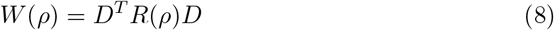

in which 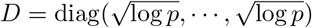 and 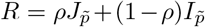 for some 0 ≤ *ρ* < 1. Here, 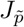 is the 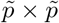 matrix with all the entries being 1. There is a one-to-one correspondence between (*ρ, T*) and (*μ, σ*^2^), the mean and variance of total edges. The details about the formulas are given in Supplementary Materials. So for any specification of (*μ, σ*^2^), the corresponding parameter (*T, ρ*) can be easily determined which in turn can be used in BIC_*T,W*_ for model selection.

## 3. Incorporation of Prior Structure into Model Selection

### 3.1. Prior structure enhanced BIC for Gaussian graphical model

In Section 2.2, it is shown that the third term in EBIC is the squared distance between a given state **z** and the origin state **0**, or in other words, the squared distance between the given graph and the empty graph. Now let us adapt BIC_*T,W*_ (7) to accommodate the prior structure of graph. For **X** ~ *N*(***μ***, Σ), suppose that graph 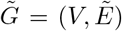 represents the prior structure (e.g., constructed based on some biological theory), and our aim is to learn the underlying true structure based on 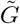 and the observations on **X**. First, we intruduce the concept of difference graph. For two graphs 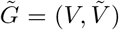 and *G* = (*V, E*), the difference graph of 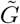 and *G* is defined as the graph which has the same vertex set *V* as 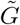 and *G* while the edge set is 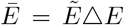. Here Δ stands for the symmetrical difference operator between two sets. Let the difference graph denoted by 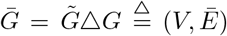. For a given prior edge set 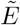, there is a one-to-one correspondence between 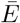 and *E*. Equivalently, there is a one-to-one correspondence between their edge vector, 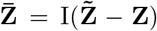 and **Z**. Replacing **z** in the third term of BIC_*T,W*_ with 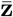, we obtain the following structural Bayesian information criterion (SBIC),

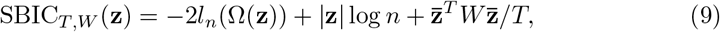

in which the first term measures the fitness between model and data, the second term measures the model complexity and the third term measures the deviation of the model from the prior structure. Minimization of (9) will lead to solutions that achieve balance between these terms. Essentially, we have assumed that 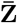 in (9) has Boltzmann distribution,

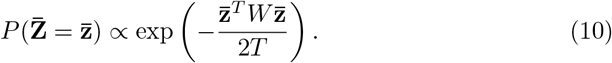

If we set *W* = diag(log*p*, ···, log *p*) just like EBIC, then (9) reduces to

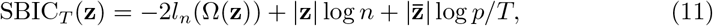

which will be used in the numerical studies in Section 5 and 6.

#### Remarks

(i) If 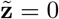, i.e., the prior structure is empty graph, then SBIC in (11) reduces to EBIC in (3), i.e., EBIC is a special case of SBIC. (ii) If *T* is large enough, then the model selected by SBIC is the same as that selected by standard BIC; if *T* is small enough, then the model selected by SBIC is just the prior graph. For other *T*, the model selected by SBIC will be a compromise of these two extreme cases. (iii) The choice of *T* in (11) relies on the expected error rate of prior structure. The expected error rate is defined as 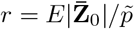, where 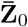 is the edge vector of the difference graph between true and prior structure. It is straightforward to show that 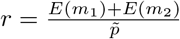, where *m*_1_ is the number of true edges that have been omitted by prior structure while *m*_2_ is the number of edges that have been mistakenly included in the prior structure. In many cases, r is more intuitive than *T*. On the other hand, we have from (10), 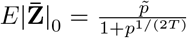, from which we have 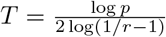. Given such one-to-one correspondence between *T* and *r*, the intuitive interpretation of *r* can guide us to determine the appropriate value for *T*. Particularly, if *r* = 0.5, then *T* = ∞, which yields the standard BIC. (iv) The knowledge about the expected error rate *r* may be derived from domain knowledge, as we did for the metabolite data in Section 6. In case that such information is not available, Bogdan et al [4, 5, 6] recommended a tuning parameter in the context of regression model with which the family wise error rate (FWER) is shown to be approximately 8% for the dataset with the sample size *n* ≥ 200 and the number of variables *p* ≥ 30. Similar strategy can also be employed in the context of GGM model to control the FWER when no prior information about r is available.

The generalization of (11) is possible. For example, it has been implicitly assumed that the probability of adding an edge to the prior graph, *p*_1_, and the probability of removing an edge from the prior graph, *p*_2_, is equal. In some cases, compared with pruning edges, we may be more inclined to add edges to the prior graph, i.e., *p*_1_ > *p*_2_. The following simple generalization of (11) can accommodate such situation,

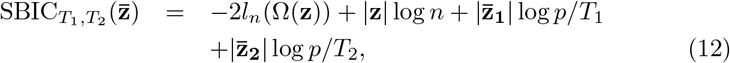

where 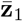 is the indicator vector of whether the entries of 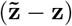 are 1, while 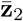 is the indicator vector for −1. Different combination of *T*_1_ and *T*_2_ reflects our different belief about the prior structure. For example, a small *T*_1_ and a large *T*_2_ indicate that we have higher confidence about the edges than the non-edges in prior structure. The cost of such flexibility is that we have to specify the values for both *T*_1_ and *T*_2_ which may be challenging in some circumstances.

As has been mentioned in Section 1, there are already several different ways to model the prior structure of network in Bayesian methods, see, e.g., Mohammadi and Wit [52], Mohammadi [53], Mukherjee and Speed [56]. In particular, the concordance function proposed in Mukherjee and Speed [56] plays the similar role as the energy function (5) in this paper. They demonstrated how different type of prior structures can be integrated into the prior distribution of the network. The network structure can then be inferred based on the samples from the posterior distribution. Note that the size of the model space for network grows super-exponentially as the number of vertices increases. Given this fact, Leday and Richardson [43] pointed out that the MCMC-based strategy for model selection of network can hardly get sufficient qualified samples that can represent the real posterior distribution, which will eventually compromise the reliability of the final estimates of network. The proposed model selection criterion (9) combined with the candidate model pool detailed in the next section provides an alternative yet feasible way to deal with these problems and demonstrates its superiority in the simulation studies compared with Bayesian network selection.

### 3.2. Construction of model pool based on prior structure

With *p* variables, the size of the model space for graphical model is 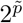. If *p* = 10, then the size of model space is 2^45^. So as *p* increases, it quickly becomes intractable to find the optimal model by searching the whole model space. There are two ways in literature to deal with this problem. One is to use the heuristic algorithm, including greedy or stepwise forward/backward search, to find the optimum of score function (Chickering et al [16]). The other is to select a subset of the model space as the candidate model pool and then use the model in this model pool which minimize the score function as our selected model (Friedman et al [22]). In this paper, we focus on the second method. A common practice for the construction of candidate model pool for graphical model is to use the solution path of graphical lasso (Friedman et al [22]). The disadvantage of graphical lasso is that it does not integrate prior structure when building the model pool. Even with SBIC in hand, we may still end up with a poor model choice. It is necessary to incorporate the prior structure into the construction of model pool. For example, in addition to the solution path of graphical lasso, we may simply include random samples from the Boltzmann distribution (10) corresponding to the prior structure as a part of the model pool. This method turns out to be inefficient for high-dimensional model given the huge size of model space. Alternatively, we can carefully devise the penalty term in graphical lasso so that the solution path can relate to the prior structure automatically. This method usually involves complex optimization algorithm that can not be easily solved based on the existing software. In the following, we propose an intuitive algorithm to build the model pool based on the prior structure, which can be easily implemented using the popular R package such as *glasso* (Friedman et al [22]) or *glmnet* (Friedman et al [24]). This algorithm bears some similarities to the well-known greedy equivalence search (GES) algorithm while the latter aims to learn the structure of directed acyclic graphical model (Chickering et al [16], Ramsey et al [59]). Recall that GES algorithm optimizes the given score function by an edge addition or removal in each step until the algorithm converges. If the true model is decomposable, it is proved that GES algorithm can consistently select the true model as sample size tends to infinity (Chickering et al [16]). Though our algorithm also involves edge addition and removal, the present objective is to construct the model pool instead of searching the optimum of score function. Specifically, the algorithm consists of the following two steps.

Step 1 (Edge enrichment): Since some edges may have been omitted by the prior graph, in this step we consider how to pick up the omitted edges. To this end, for a given increasing sequence of 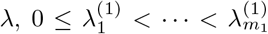, we solve the following optimization problems for *i* = 1, ···, *m*_1_,

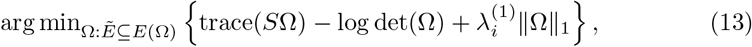

where, *E*(Ω) is the edge set of graph corresponding to Ω, 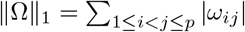. In (13) we fix the edges in prior structure and consider how to enrich it by selecting the edges from the rest edges. This step leaves us *m*_1_ graphs denoted by *G*^(*i*)^ for *i* = 1, ···, *m*_1_ respectively.

Step 2 (Edge pruning): Each *G*^(*i*)^ (*i* = 1, ···, *m*_1_) from the first step contains the prior structure. Since some redundant edges may have been mistakenly included in the prior structure, we aim to prune these edges from these graphs in this step. To this end, for a given *G*^(*i*)^ (1 ≤ *i* ≤ *m*_1_), we solve the following (2) (2) *m*_2_ optimization problems for an increasing sequence 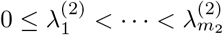,

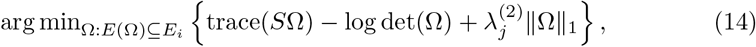

where *E_i_* is the edge set of graph *G*^(*i*)^. In (14), for a given graphical model *G*^(*i*)^, we consider how to prune the false edges that have been included in the prior structure. This step leaves us graphs *G*^(*ij*)^ for *i* = 1, ·, *m*_1_ and *j* = 1, ···, *m*_2_. Thus there are total *m*_1_*m*_2_ candidate models in the final model pool.

#### Remarks

(1) If the prior structure is an empty graph, then only *edge enrichment* step is involved to build the model pool; if the prior structure is a complete graph, then only *edge pruning* step is involved. The model pools for these two extreme cases turn out to be the same as that from the standard lasso. (2) The two-step algorithm above is implemented through the graphical lasso (Friedman et al [23]); nevertheless, the algorithm can also be equivalently implemented through the neighborhood method (Meinshansen and Buhlmann [49]). The consistency of neighborhood method is guaranteed by the property of lasso for high-dimensional regression model (Zhao and Yu [87]). When neighborhood method is used, the R package *glmnet* (Friedman et al [24]) can be employed to facilitate the computation.

A concern raised by the reviewers is about the performance of the proposed algorithm when there exists big discrepancy between the prior and true network. Firstly, we note that the proposed two-step algorithm always includes the solution path of lasso as a part of the model pool; consequently, the model pool can cover the true model if the sample size n is reasonably large with respect to the number of vertices p. As far as SBIC is concerned, this issue essentially is about how to select the temperature parameter *T*. If we have a good estimate of the discrepancy between prior and true network, or equivalently the average error rate *r*, then an appropriate *T* can be selected. In that case, a bad prior network will make the last term in SBIC relatively small and the model selection will be mainly determined by other terms in SBIC. However, if we mistakenly assume a small r for an actually large discrepancy, that will have a negative effect on the model selection. In that situation, sensitivity analysis of the result with respect to the tuning parameter *T* is recommended, based on which tuning parameter can be chosen to ensure a robust network selection, see section 5.1 for details.

## 4. Consistency of SBIC

In this section we investigate the theoretical properties of SBIC. It is proved that under the given assumptions SBIC can consistently select the underlying model for high-dimensional Gaussian graphical model, where the number of vertices may increase as sample size increases.

First let us introduce some notations for ease of exposition. Recall that **z** is the 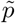-dimensional edge vector indicating whether or not there is an edge between two given vertices. Define 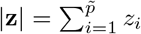 and let **z**_0_ be the edge vector corresponding to the true graph *E*_0_ under consideration. Let 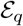 denote the graph set with no more than *q* edges and 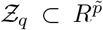 the corresponding edge vector set. Let 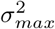 denote the largest diagonal component of the true covariance matrix Σ_0_ while *λ_max_* denote the largest eigenvalue of true precision matrix Θ_0_. For a given positive semi-definite matrix *W*, let *τ*_max_ and *τ*_min_ be the largest and smallest eigenvalue of *W* respectively. With these notations in hand, the consistency for BIC_*T,W*_ (7) and SBIC (11) are proved in Theorem 1 and 2 respectively. For BIC_*T,W*_, the following assumptions are involved.

**Assumption 1**. 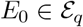 is decomposable;

**Assumption 2.** *p* = *O*(*n^κ^*) for some 0 < κ < 1;

**Assumption 3**. ∃ constant *C* > 0 such that 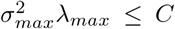 and 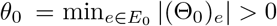

**Assumption 4.** ∃ϵ > 0 such that 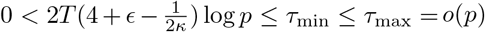.

### Theorem 1.

Under Assumptions 1-4, the model selection procedure based on BIC_*T,W*_ given in (7) is consistent, i.e., as *n* → ∞ we have

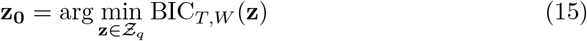

in probability.

Now let us consider SBIC (11) in which prior structure is available for the underlying graphical model. Recall that 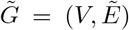 is the prior structure, 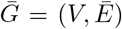 is the difference graph between 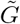 and *G* and 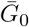 is the difference graph of prior graph 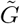 and true graph *G*_0_. Here we have assumed 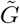 and *G*_0_ have the same vertex set. In order to prove the consistency of SBIC (11, Assumptions 1 and 4 have to be replaced by the following Assumptions 1 and 4′.

**Assumption 1** 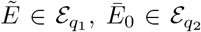 for some integers *q*_1_ and *q*_2_ and *E*_0_ is decomposable.

**Assumption 4** For 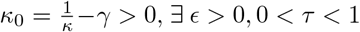, ∃ϵ > 0, 0 < *τ* < 1 such that *τκ*_0_ > 4+*ϵ*.

Assumption 1 says that 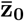 has at most *q*_2_ nonzero components which means that we can reach the true edge set *E*_0_ by adding or deleting at most *q*_2_ edges from the prior edge set 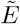 and so 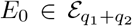. Given the observations 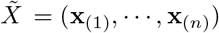, we have the following result.

### Theorem 2.

Under Assumption 1′ and 2, 3 and 4′, SBIC (11) can consistently select the true graph structure *G*_0_, i.e., as *n* → ∞, we have

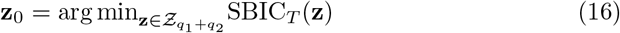

in probability.

The details of the proof for Theorem 1 and 2 are provided in Supplementary Materials. It should be noted that, in order to facilitate the proof, we have imposed strong assumptions on the dimensionality and the underlying graphical structure such as decomposability. It is possible that the results still hold if some of these assumptions are relaxed. In particular, both the evaluation of SBIC and implementation of two-step algorithm do not depend on the decomposability of the underlying graph.

It also should be pointed out that we only discussed the problem of model selection for GGM in this study and did not investigate the problem of its statistical inference. It is implicitly assumed that the statistical inference can be reasonably conducted after the model is selected, though this is not always the case in practice. Additional assumptions including irrepresentability and beta-min condition have been suggested in literature to ensure the consistency of such estimate (B*ü*hlmann et al [9]). Recently, several novel methods that integrated the model selection and the statistical inference have been proposed for GGM. For example, Ren et al [63] employed a two-dimensional regression model to estimate each entry of precision matrix and the asymptotical distribution is shown to be normal. Jankova and van de Geer [33] adapted the neighborhood method of Meinshansen and Buhlmann [49] based on the Karush-Kuhn-Tucker (KKT) conditions and proposed an estimator for each entry of precision matrix which is shown to converge asymptotically to the normal distribution. R package *SILGGM* has been developed to implement the statistical inference for GGM using these new methods (Zhang et al [84]).

## 5. Simulation Studies

In this section the proposed algorithm is evaluated based on simulated data. In the first example, we consider a tree graph. For tree graphs, the well-known Chow-Liu algorithm can optimally and efficiently learn the structure (Chow and Liu [17], Edwards et al [21], Kirshner et al [38]). We will compare the performance of our method with Chow-Liu algorithm. In addition, there is a temperature parameter *T* involved in SBIC. Though ideally *T* should be determined based on the prior information, e.g., expected error rate, in practice prior information are often biased to some extent. So we also perform sensitivity analyses to evaluate the robustness of our method with respect to the misspecification of *T*. In the second example, we go beyond the tree model and consider the randomly generated graphs which may be non-decomposable. Based on the simulated data from these graphs, we compare the proposed algorithm with other popular model selection methods in literature. It is demonstrated that the proposed algorithm can outperform these existing algorithms under the given scenarios in terms of two indices, true positive rate (TPR) and false positive rate (FPR), which are defined as

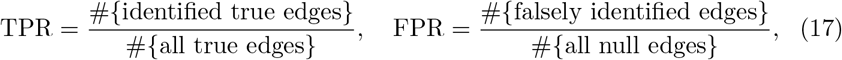

for which a higher TPR and lower FPR indicate a better model selection.

### 5.1. Sensitivity analysis based on tree model

Consider a Gaussian graphical model with fixed tree structure. Specifically, let *X* = (*X*_1_, ···, *X*_40_) be a random vector with *X*_1_ ~ *N*(0,1). For *i* = 2, 3, 4, we have *X_i_* = *αX*_1_ + *ϵ_i_* with *ϵ_i_* ~ *N*(0,1). For *i* = 5, 6, 7, we have *X_i_* = *αX*_2_ + *ϵ_i_* with *ε_i_* ~ *N*(0,1). For *i* = 8, 9, 10, we have *X_i_* = *αX*_3_ + *ϵ_i_* with *ϵ_i_* ~ *N*(0,1). In the same fashion, all the variables can be generated. The structure of *X* is shown in the right plot in Figure 1. The left graph in Figure 1 is used as the prior structure.

**Figure 1:**
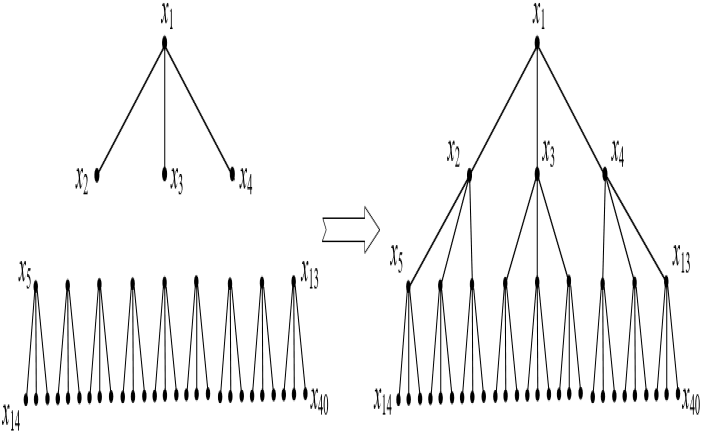
The graphs involved in Section 5.1. The left one is used as the prior graph while the right one is the true graph.

Two plots in Figure 2 present TPR and FPR as the function of *α* respectively. In each plot two curves are drawn in which the solid curve corresponds to the model pool constructed from standard lasso while the dashed curve corresponds to the model pool constructed from two-step algorithm. In both cases SBIC is employed to select the model in which temperature parameter is set based on true error rate *r* = 9/780. Here the replication is *N* = 100; the sample size is *n* = 60. The difference in each plot reflects the difference between the two model pools. From Figure 2 it is obvious that TPR from two-step algorithm is higher than that from standard lasso while FPR from to-step algorithm is lower than that from standard lasso. In particular, the difference becomes more prominent when the association among the variables is weak.

**Figure 2:**
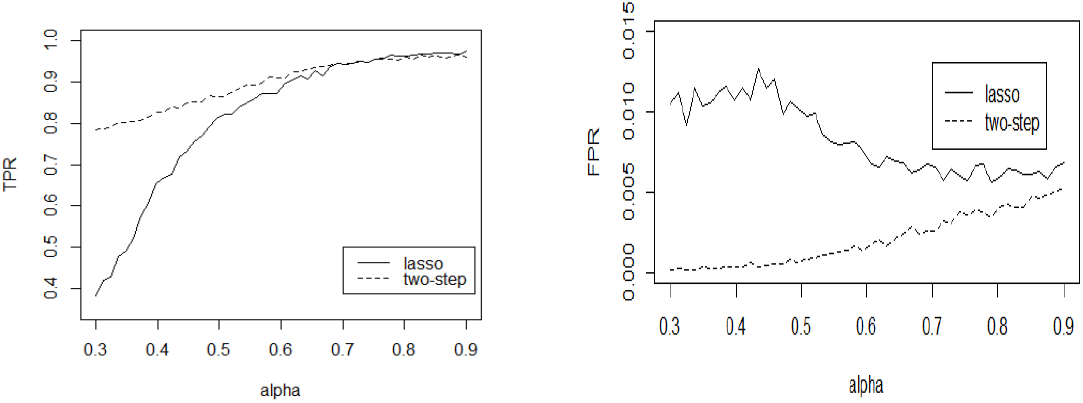
The left plot is the TPR versus association strength *α* and the right plot is the FPR versus *α*. The solid lines correspond to the model pool constructed based on lasso while the dashed lines correspond to the model pool constructed based on two-step algorithm.

Table 1 lists the results for multiple specifications of *T* and Chow-Liu algorithm under different scenarios. Specifically, the sample sizes are *n* = 50, 100; the number of replication is *N* = 100. Three choices of association strength are *α* = 0.3, 0.4, 0.5. As for temperature parameter T, five choices for expected error rate, *r* = 3/780, 9/780,18/780, 27/780, 36/780, are considered, from which *T* can be derived based on the formula in Section 3.1. For example, the value of *T* in SBIC_1_ corresponding to *r* = 3/780 can be shown to be 0.332.

**Table 1:** Sensitivity analysis of structural Bayesian information criterion (SBIC) with respect to the temperature parameter. The first number in the parentheses is true positive rate (TPR), and the second number is false positive rate (FPR). The results for BIC and Chow-Liu algorithm are also listed. The number of replication for each result is *N* = 100.

|  | n=50 |  |  |  |  |  | n=100 |  |  |  |  |  |
| --- | --- | --- | --- | --- | --- | --- | --- | --- | --- | --- | --- | --- |
| | $\alpha = 0.4$ | | $\alpha = 0.5$ | | $\alpha = 0.6$ | | $\alpha = 0.4$ | | $\alpha = 0.5$ | | $\alpha = 0.6$ | |
| SBIC<sub>1</sub>+TS | (0.794, 0.0000) |  | (0.838, 0.0003) |  | (0.876, 0.0009) |  | (0.894, 0.0005) |  | (0.941, 0.0007) |  | (0.971, 0.0009) |  |
| SBIC<sub>2</sub>+TS | (0.809, 0.0003) |  | (0.844, 0.0008) |  | (0.885, 0.0012) |  | (0.912, 0.0008) |  | (0.953, 0.0007) |  | (0.974, 0.0009) |  |
| SBIC<sub>3</sub>+TS | (0.816, 0.0008) |  | (0.848, 0.0009) |  | (0.894, 0.0015) |  | (0.902, 0.0008) |  | (0.952, 0.0008) |  | (0.982, 0.0013) |  |
| SBIC<sub>4</sub>+TS | (0.828, 0.0008) |  | (0.870, 0.0014) |  | (0.899, 0.0019) |  | (0.904, 0.0008) |  | (0.949, 0.0007) |  | (0.975, 0.0012) |  |
| SBIC<sub>5</sub>+TS | (0.827, 0.0011) |  | (0.870, 0.0015) |  | (0.902, 0.0026) |  | (0.902, 0.0007) |  | (0.951, 0.0010) |  | (0.979, 0.0012) |  |
| BIC+TS | (0.916, 0.0323) |  | (0.948, 0.0283) |  | (0.958, 0.0198) |  | (0.972, 0.0171) |  | (0.991, 0.0146) |  | (0.996, 0.0113) |  |
| BIC+Lasso | (0.664, 0.0275) |  | (0.836, 0.0261) |  | (0.907, 0.0261) |  | (0.932, 0.0282) |  | (0.981, 0.0240) |  | (0.993, 0.0134) |  |
| Chow-Liu | (0.653, 0.0179) |  | (0.839, 0.0079) |  | (0.917, 0.0039) |  | (0.9107, 0.0050) |  | (0.979, 0.0009) |  | (0.991, 0.0003) |  |

From Table 1 it can be seen that: (1) For the combination of SBIC and two-step algorithm, in most cases both TPR and FPR increase if *T* increases. For the rows with the error rate other than the true value *r* = 9/780, the moderate deviation of r from the true value does not have big impact on the final results; (2) The combination of BIC and lasso yields large false positive rates which explains the popularity of EBIC; the combination of BIC and two-step algorithm yields better results, i.e., higher TPR and lower FPR, especially when the association strength *α* is low and sample size is small; (3) In most cases the combination of SBIC and two-steep algorithm yields higher TPR and lower FPR than Chow-Liu algorithm. Nevertheless, compared to the combination of BIC and lasso, Chow-Liu algorithm yields comparable TPR and lower FPR.

In summary, if prior structure is available for graphical model, then both model selection criterion and candidate model should incorporate such information. The results from the proposed procedure also demonstrate robustness to the misspecification of temperature parameter.

### 5.2. Comparison with other model selection strategies based on general graphical model

In this section, we extend the tree graph considered in Section 5.1 to the randomly generated graph with up to 200 vertices. The proposed model selection algorithm is compared with four popular model selection methods in literature, including (1) Bayesian method; (2) CV(cross validation)+Lasso; (3) EBIC+Lasso; (4) BIC+rLasso. For Bayesian method, independent edge inclusion indicator variables are assumed for the edges set. The same Bernoulli prior is assumed for all indicator variables. With a given network structure, we then use the G-Wishart distribution as the prior distribution of precision matrix, which is known to be conjugate distribution for the normally distributed data. For second method CV+Lasso, model selection criterion is CV while model pool is built by lasso. No prior information is involved here. For EBIC+Lasso, model selection criterion is EBIC while model pool is built by lasso. Prior structure is used in EBIC in the same way as SBIC; however, the tuning parameter in (3) is set to be fixed at λ = 0.5 as commonly suggested in literature. Method 4 is proposed in Ma et al [48] where rLasso stands for residual lasso. Prior information is used in rLasso to construct the model pool. Specifically, when a part of the structure is known a priori with certainty, Ma et al [48] proposed to use lasso to construct the model pool based on the residuals from the linear regression of each variable on its known neighbors. Given such model pool, they then employed BIC to select graphical model. Such method to build the model pool, however, will be biased when the prior structure involves randomness. The two-step algorithm proposed in this paper takes all the vertices into consideration in the enrichment step which theoretically leads to a less biased model pool than that from rLasso. Furthermore, since the model pool can not fully reflect the randomness information in prior structure, the combination of SBIC and the proposed model pool should have better performance if the prior information have been reasonably specified.

Specifically, we first randomly generate a *p* × *p* adjacency matrix *M*_1_ as the true structure, in which the number of edges, i.e., the number of 1’s among the off-diagonal entries of *M*_1_, is set to be 100. The adjacency matrix of prior structure *M*_2_ is generated by randomly changing 100α percent of the 1’s entries of *M*_1_ to 0 and the same number of 0’s entries to 1. Given *M*_1_, a symmetrical matrix with 1’s on its diagonal is generated which has the same edge set as *M*_1_. Each of nonzero entries in this matrix is generated from *N*(0,1) distribution. By tuning the diagonal element 1 to some value *β*, we can always get a positive definite matrix *K*, which will be used as the precision matrix in this study. Here we choose *β* = 1.1 – λ_min_ (A) in which λ_min_ (A) denotes the minimum eigenvalue of matrix *A*. For *p* = 100, 200, *α* = 0.7, 0.5,0.3, 0.1, and sample size *n* = 100, 200, Table 2 lists the results of TPR and FPR in each scenario for all the five model selection strategies. For Bayesian method, the inclusion probability for each edge is set to be *θ* = 200/*p*(*p* – 1), which means that we do not assume any specific structure for the graph in its prior distribution other than the sparsity. For the prior G-Wishart distribution of precision matrix, *W_G_*(*b, D*), we set *b* = 3 and *D* the *p* × *p* identity matrix. The burn-in for sampling from the posterior distribution is set to be 5000. For a given vertex pair (*X_i_, X_j_*), if 50% precision matrix samples have nonzero (*i, j*)th entries, then we define an edge between *X_i_* and *X_j_*; Otherwise, no edge is defined between *X_i_* and *X_j_*. As for methods 2 to 5, the number of tuning parameters to build the model pool is set to be 100. The temperature parameter in SBIC is set based on the discrepancy rate between real and prior networks, i.e., *α* = 0.7, 0.5,0.3, 0.1. Note since method CV+Lasso does not involve any prior information, so it has the same TPR’s and FPR’s in all the four discrepancy situations in Table 2. It can be seen that the performance of Bayesian method is sensitive to the prior information. It should be noted that Bayesian method, which is implemented through the R package *BDgraph* (Mohammadi and Wit [52]), is much more time-consuming than the other four methods; CV+Lasso tends to select the graphs with too many false edges; EBIC+Lasso tends to omit too many true edges. Though BIC+rLasso has a better performance than CV+Lasso and EBIC+Lasso, in most cases, it still has a high probability to omit the true edges and select the false edges. As for our proposed strategy, it works well and reaches a good balance between TPR and FPR, and compared to other four methods, yields higher TPR while lower FPR in most cases, especially when the discrepancy between prior structure and true structure becomes smaller.

**Figure 3:**
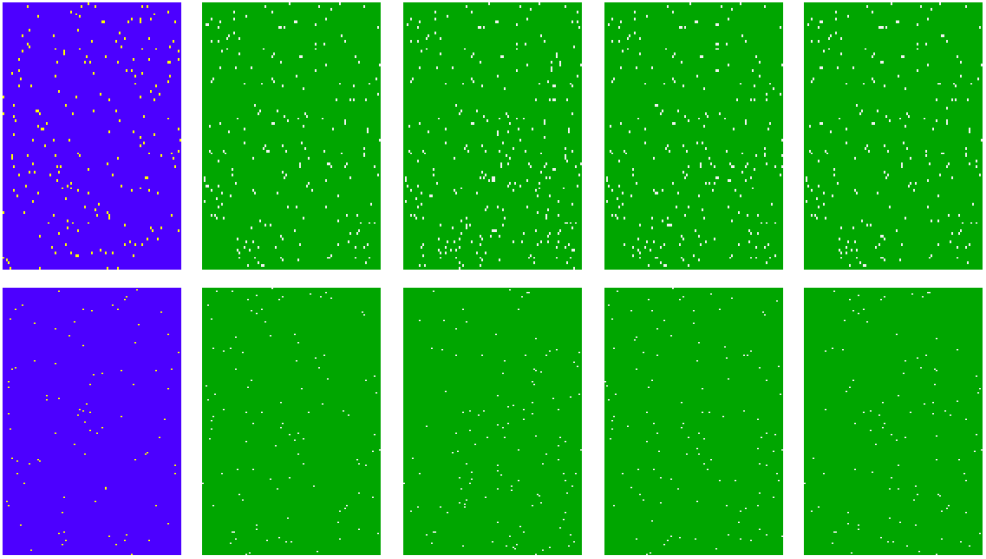
A realization of the real and prior networks in simulation studies in Section 5.2. Dots indicate the nonzero conditional associations between vertices. The five plots in the first row are for network with *p* = 100 vertices. The first one is the true network, and the discrepancy between the other four networks and true network are, from left to right, *α* = 70%, 50%, 30%, 10% respectively. These networks are used as prior networks respectively. Similar explanation applies to the networks in second row which have p = 200 vertices.

**Table 2:**
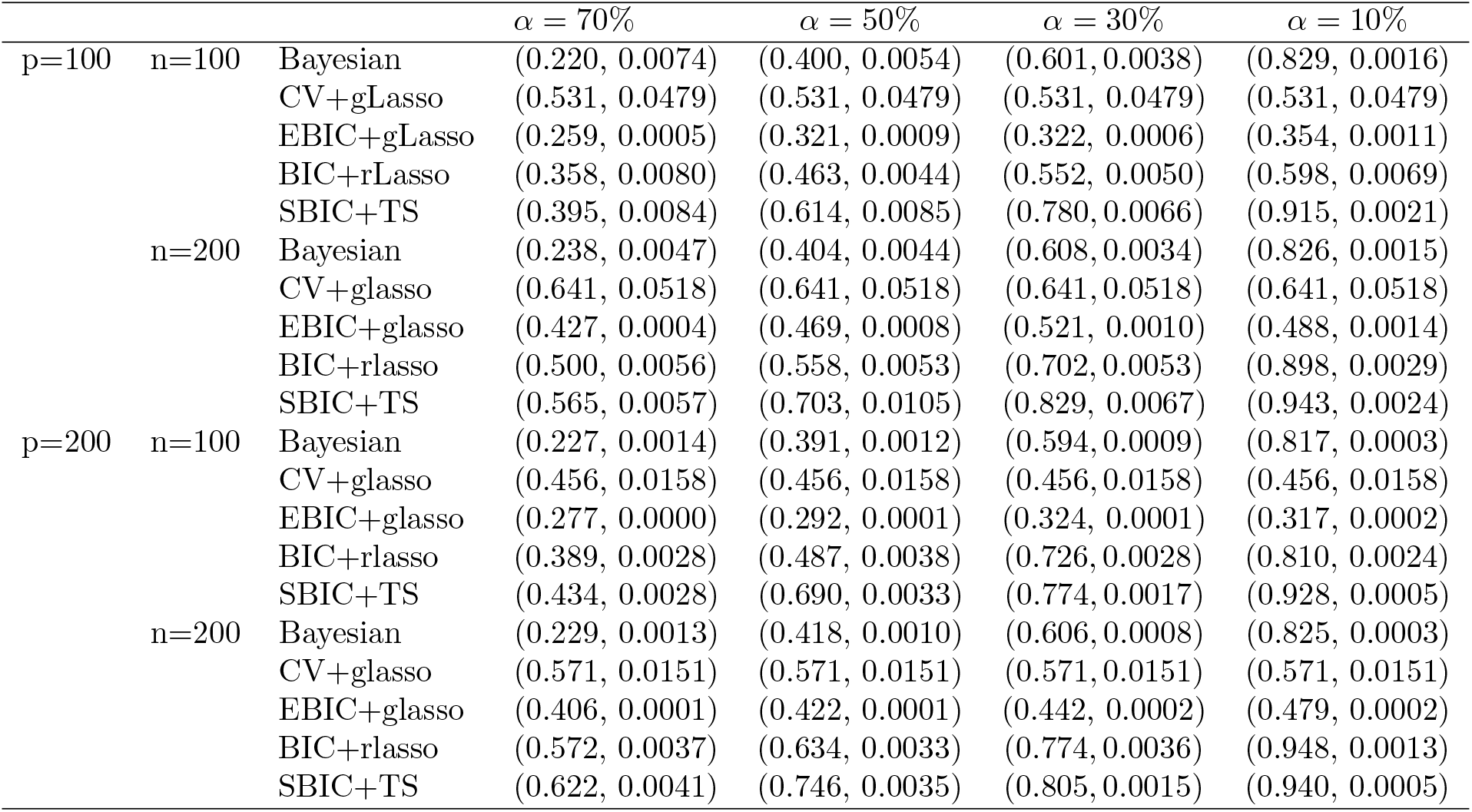
Performance comparison of different model selection strategies. The first number in the parentheses is true positive rate (TPR), and the second number is false positive rate (FPR). Number α stands for the proportion of the edges that have been falsely included in prior network.The number of replication for each result is *N* = 100.

## 6. Metabolite Network in Human Gut

Metabolites in human body are intrinsically related with different diseases. Understanding the relationship between metabolites are helpful to design appropriate treatment. To this end, multiple methods have been proposed in literature to identify the structure of metabolite networks. For example, Gao et al [27], Karnovsky et al [40] used the biochemical domain knowledge to construct the metabolite network. Barupal et al [3], Grapov et al [29] constructed the network based on structural similarity and mass spectral similarity of metabolites. The metabolite prior networks in this paper are constructed using the similar method to that in Gao et al [27], Karnovsky et al [40].

The dataset involved comes from the New Hampshire Birth Cohort Study, an ongoing prospective cohort study of women and their young children Madan et al [46]. The dataset was obtained from metabolomics characterizations of stool samples collected from infants at approximately six weeks to one year of age. Sample preparation (with some modifications), ^1^H NMR data acquisition, and metabolites profiling procedures have been previously described in Banerjee et al [2], Brim et al [8], Pathmasiri et al [57], Sumner et al [73, 74]. Chenomx NMR Suite 8.4 Professional software (Edmonton, Alberta, Canada) was used to determine relative concentration (Weljie et al [82]) of selected metabolites from a curation of list of metabolites that are associated with host-microbiome metabolism, see Li et al [44], Paul et al [58]. This resulted in a total of 882 observations for 36 metabolites in this data set. All the observations for metabolites were standardized so that they have zero mean and unit standard error, see van den Berg [78]. In the following, we consider how to learn the structure of the network among these metabolites using the algorithm proposed in Section 3.

Specifically, we use pathway analysis to construct the prior structure. These pathway information are obtained from biological database Kyoto Encyclopedia of Genes and Genomes (KEGG) which provides state-of-the-art information about the metabolites and their pathways. Note that each of the targeted metabolites is listed with its associated KEGG Compound ID. Compound information for small molecules in the KEGG database can be retrieved using KEGGREST, a client API written for R (Dan Tenenbaum (2018). KEGGREST: Client-side REST access to KEGG. R package version 1.22.0). Using functions in the KEGGREST library, the database resource was queried in the R language to retrieve the list of one or more pathways associated with each metabolite. With the pathway information in hand, for two given metabolites *X_i_* and *X_j_*, let the pathways associated with *X_i_* and *X_j_* be *Z_i_* = {*Z*_*i*1_, ···, *Z_im_i__*} and *Z_j_* = {*Z*_*j*1_, ···, *Z_jm_j__*} respectively. Denote the common pathways of *X_i_* and *X_j_* by *Z_ij_* = *Z_i_* ⋂ *Z_j_* and let

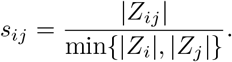

If *s_ij_* ≥ 0.8, then we define an edge between *X_i_* and *X_j_* in the prior graph. With threshold equal to 0.8, there are 27 edges in the prior graph. With threshold equal to 0.6, there are 117 edges in the prior graph. We use the difference of these two number as the expected number of edges in difference graph between the prior and true network which in turn implies that the value of temperature parameter involved in SBIC is *T* = 1. As for the construction of model pool, we set *m*_1_ = *m*_2_ = 200 with λ_max_/λ_min_ = 0.01 in (13) and (14), where λ_max_ represents the minimal λ at which the graph has no edge. Then based on SBIC (11) and two-step algorithm, we obtain the final network. Comparison of the prior and the final network reveals that there are 153 edges added to and 3 edges removed from the prior network. Figure 4 shows the added edges. The three removed edges are (Methionine, Tryptophan), (Glutamate, Histidine), (Asparagine, Valine) respectively.

**Figure 4:**
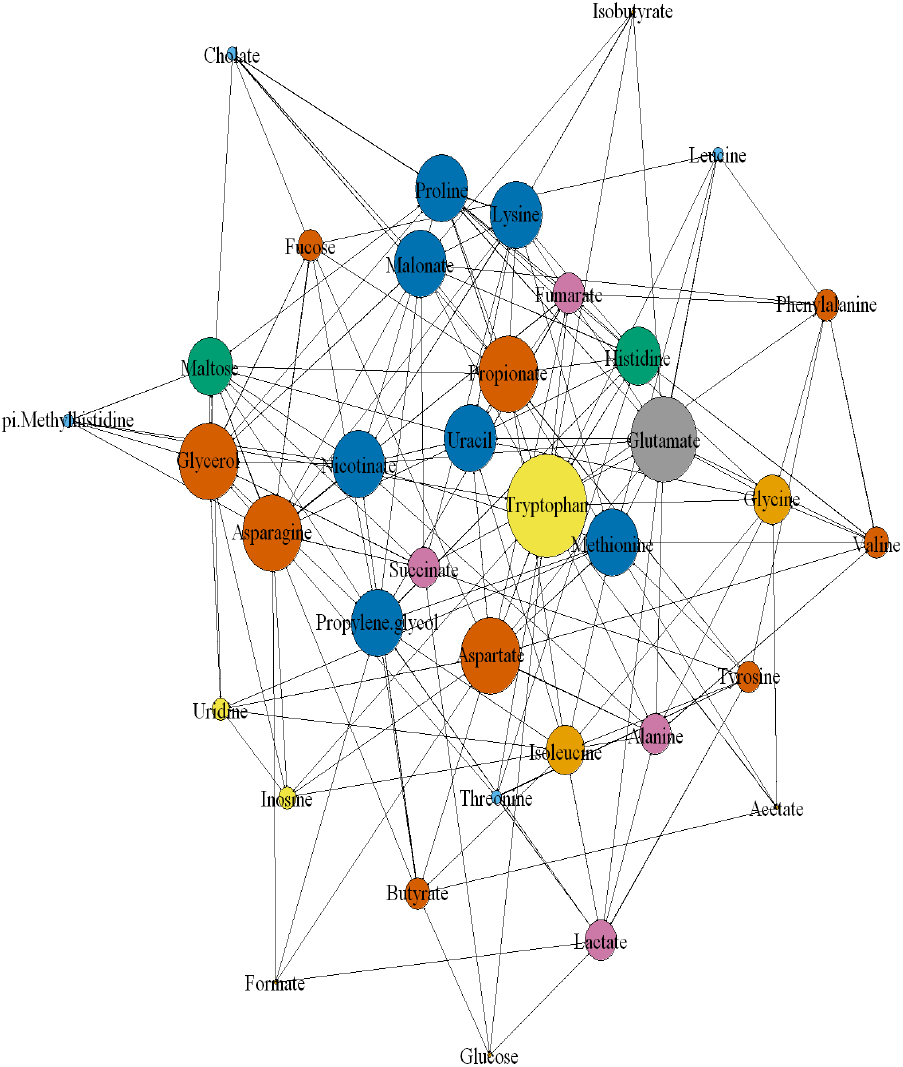
The edges that appear in the final network while have not been included in prior network.

A primary question here is that whether the edges that are defined by pathway reflect the association between metabolites. If a pathway does not contain any information about association between metabolites, then such prior network can be regarded as built just randomly. Then the probability *p*_1_ that an edge is removed from and the probability *p*_2_ that an edge is added to the prior network should be equal. Thus we can consider the following hypothesis testing problem, H_0_: *p*_1_ = *p*_2_. The test statistic involved is 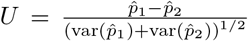 where 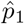 and 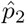 are the maximum likelihood of *p*_1_ and *p*_2_ respectively. In light of central limit theorem, it can be shown that the p-value for the hypothesis above is 0.0234. With such a p-value, we can tentatively assert that pathways have statistically significant effect on the association between metabolites.

One potential concern about the previous analysis is that the conclusion may be biased by the prior structure. However, we still can use the following method to validate this conclusion. Specifically, we just consider the added edges in Figure 4 which are not involved in prior structure. For any given 0 < *s* < 0.8, we construct the prior network *E_s_* by using the same procedure as above, i.e, add an edge for (*X_i_, X_j_*) if *s_ij_* ≥ *s* otherwise not. Note for *s* = 0.8 there are 153 added edges among total 603 edges, apart from the 27 prior edges, that have been selected by the proposed method. Imagine that if pathways have no impact on the association of metabolites, then the proportion of 153 added edges in *E_s_* should be the same as for *s* = 0.8, i.e., 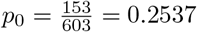. Define *p_s_* the probability of the edges in Figure 4 falling into *E_s_*, then the null hypothesis is H_0_: *p_s_* = *p*_0_. For *s* = 0.1, 0.2, ···, 0.6, the estimate 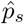 can be shown to be (0.2578,0.2596,0.2800, 0.3300, 0.3630,0.4111) and the corresponding p-value for the hypothesis H_0_ are (0.3959,0.3658,0.1287,0.0059, 0.0011,0.0003). Based on these results, we can say that pathway does contain the association information between metabolites. The more pathways two metabolites share, the more likely their concentrations are related. Figure 5 shows the empirical probability of nonzero association as a function of threshold.

**Figure 5:**
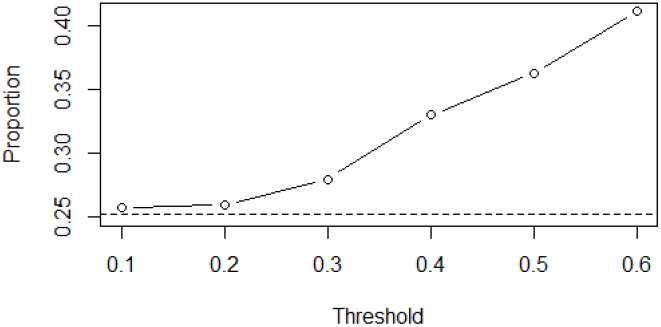
The proportion of the added edges in prior network *E_s_* as a function of threshold *s*. The bottom dashed line corresponds to the line under the null hypothesis.

It should be stressed that the discussion above does not mean that prior network must have to share some common information with the data. If a prior network is theoretically sound, such prior network is also feasible. However, if a prior network can find the support from both the theory and data, in our view, it is more advantageous than the one with support just from theory or subjective belief.

We have confirmed that part of the association among metabolites can be attributed to pathway. The next question we aim to address is that whether all the association among metabolites can be explained solely by pathway. To try to answer this question, first we define a densest prior structure among metabolites based on pathway. Specifically, whenever two metabolites have any pathway in common, we define an edge between them and no edge otherwise. By comparing the network in Figure 4 to this prior structure, we found that there are 20 edges which are not covered by the prior structure. In other words, pathway cannot explain all the association between metabolites.

These 20 edges are listed in Table 3. Among these 20 edges, 8 edges are related with malonate, 8 edges are related with propylene glycol and the rest are related with π-Methylhistidine. Malonate is a well-known competitive inhibitor of succinate dehydrogenase (SDH) while SDH is a complex of four polypeptides (SDH A–D) that catalyzes the conversion of succinate to fumarate and functions in mitochondrial energy generation, oxygen sensing and tumor suppression. Propylene glycol is a widely used drug vehicle with serious side effects reported in clinical studies and recognized toxicity (Morshed et al [54, 55]). In light of these existing studies, it is not surprising to find out their wide connections with other metabolites even they do not share any pathway.

**Table 3:** Edges in Figure 4 that can not be covered by conventional pathway analysis

|  |  |
| --- | --- |
| Malonate | Asparagine, Cholate, Isobutyrate,<br>Tryptophan, Phenylalanine, Propionate,<br>Succinate, Lysine |
| Propylene glycol | Butyrate, Formate, Methionine,<br>Fumarate, Histidine, Isoleucine,<br>Maltose, Fucose |
| π-Methylhistidine | Asparagine, Maltose, Nicotinate,<br>Tryptophan |

In summary, metabolic pathway can explain part of the association between the metabolites but not completely. This may be explained by the fact that conventional metabolic pathway datasets only focus on the endogenous reactions occurring within the cell. It is possible that some important reactions may be omitted by conventional pathway analysis. However, by appropriately combining prior knowledge with empirical data analysis, the proposed algorithm can discover these omitted associations efficiently.

## 7. Conclusion

We develop a novel method to select the high-dimensional Gaussian graphical model with the aid of prior structure. Such prior structure is often the result of biological knowledge. The algorithm consists of two parts. In the first part, we propose a novel model selection criterion called structural BIC which is a generalization of extended BIC. In second part, we propose a two-step algorithm to construct the candidate model pool which incorporates the prior structure into the candidate model through edge enrichment and pruning. It is proved that under some mild conditions the structural BIC is a consistent criterion for graphical model selection. Simulation results validate the efficacy and robustness of the algorithm.

We apply the proposed algorithm to the concentration data of metabolite in human gut for which the prior network is constructed through the pathways shared by metabolites. It is shown that pathway is a statistically significant factor for the association between metabolites. As the network based on the pathway analysis have been widely used in many fields, these findings provide statistical basis for such practice. We also find new relationships between metabolites that can be omitted by conventional pathway analysis.

It is possible to use other types of prior network for metabolites, e.g, the structural similarity based prior network. Other biological network such as gene regulation network or microbial interaction network can also be analyzed based on our method if the prior structure can be properly defined. The algorithm can be adapted for the binary data such as Ising model. It is known that model selection with prior structure for Ising model is complex and little work has been done in this respect. Our method provides a possible solution to this issue and deserves further investigation in the future.

## Acknowledgements

We are grateful to the participants and staff of the New Hampshire Birth Cohort Study for providing the processed metabolomics data. This work is supported in part by US National Institutes of Health grants (R01LM012012; R01LM012723; P20ES018175; P01ES022832; UG3OD023275), US Environmental Protection Agency grant (RD83459901).

## Supplementary Materials

Software in the form of R code is available at *https://github.com/hoenlab/SBIC*. The Supplementary Materials for this article are available in Appendix A-C, which include the marginal distributions of homogeneous Boltzmann distribution, the proofs of Theorem 1 and 2 and the prior graph for metabolomic data in section 5.

